# Droplet based low input proteomic platform for rare cell populations

**DOI:** 10.1101/2023.09.11.557098

**Authors:** Matthew Waas, Amanda Khoo, Pirashaanthy Tharmapalan, Curtis W. McCloskey, Meinusha Govindarajan, Bowen Zhang, Shahbaz Khan, Paul Waterhouse, Rama Khokha, Thomas Kislinger

**Affiliations:** Princess Margaret Cancer Centre, University Health Network, Toronto, Ontario, M5G 2C1, Canada; Department of Medical Biophysics, University of Toronto, Toronto, Ontario, M5G 1L7, Canada

## Abstract

Deep proteomic profiling of rare cell populations has been constrained by sample input requirements. Here, we present DROPPS, an accessible low-input platform that generates high-fidelity proteomic profiles of 100 - 2,500 cells. By applying DROPPS within the mammary epithelium, we elucidated the connection between mitochondrial activity and clonogenicity, discovering and validating CD36 as a marker of progenitor capacity in the basal cell compartment. We anticipate DROPPS will accelerate biology-driven proteomic research for a multitude of rare cell populations.

## Introduction

The mammary gland is a dynamic organ that substantially transforms during pubertal development, sex hormone triggered reproductive cycling, and pregnancy.^1–3^ The mammary epithelium that forms the branching system of ducts and alveolae constitutes 10-15% of the overall volume of the mammary gland and contains cells of heterogenous function including hormone sensing, secretory, stem, and progenitor activity.^1–3^ Stem and progenitor cells essential for mammary ductal expansion constitute < 5% of the epithelium.^4^ These rare cells are the cells-of-origin for the most aggressive, deadly subtypes of breast cancers (*i.e*., triple negative, basal-like and claudin-low subtypes).^5–7^ An attractive prevention strategy for breast cancer is to develop therapeutic modalities to monitor and target the oncogenic precursors as they arise.^8^ Though various markers have been reported to enrich stem/progenitor capacity^9–12^, there remains no consensus regarding the exact trajectory of mammary epithelial cells and how the cells enriched by distinct markers relate to each other. Improving our knowledge of the features and regulation of stem/progenitor cells has the potential to improve clinical outcomes for breast cancer patients.

In-depth proteomic interrogation of cells, tissues, and fluids is often performed with mass spectrometry (MS) workflows, where proteins are extracted and enzymatically digested to obtain peptides.^13–15^ Due to the extensive sample processing required prior to data acquisition, such proteomic methods often require 10,000s – 1,000,000s of cells (roughly, 10 – 100’s µg) as starting material, which can preclude the analysis of rare cell populations or other sample-limited systems.^16^ Recent single-cell or low-input proteomic methods typically leverage low-volumes and minimal manual manipulation to reduce losses during sample preparation.^17–19^ Despite these technical advances, many low-input methods only support specific study design (*e.g.*, isobaric labeling with booster channel) or require specialized equipment (*e.g.*, microfabrication capabilities, microfluidic cell isolation systems). To address this limitation, we have developed an accessible low-input proteomic method for studying rare populations of cells, <u>Dr</u>oplet-based <u>o</u>ne-pot <u>p</u>reparation for <u>p</u>roteomic <u>s</u>amples (DROPPS), that is simple to implement and easily integrates with fluorescence activated cell sorting (FACS). DROPPS utilizes commercially available and economical materials to enable reproducible and biologically-informative proteomic analysis of small numbers of cells (100s -1,000s of cells, or 20 ng – 1 µg) from *in vitro* and *in vivo* systems.

We previously generated proteomic profiles of FACS purified luminal and basal compartments of the mouse mammary epithelium and discovered differences in preferred metabolic pathways.^20^ Investigating the functional consequences of these differences revealed a positive relationship between mitochondrial membrane potential and clonogenicity, particularly in the basal compartment.^20^ Here, we expand upon those findings by applying DROPPS to subpopulations of mammary epithelial cells FACS purified according to mitochondrial membrane potential to investigate the proteomic features of enhanced clonogenic activity. Strikingly, in basal cells, high mitochondrial activity is associated with proliferation pathways and adhesion-dependent signaling. We identify CD36 as a marker for basal cells with high mitochondrial potential, increased fatty acid abundance, and enhanced clonogenicity. These findings further define the relationship between energy metabolism and progenitor cell capacity, providing insights into mammary basal cell compartment functional heterogeneity. Overall, we demonstrate that DROPPS is an accessible proteomics workflow to enhance and accelerate biological discovery for rare cell populations.

## Results

### Droplet digestion for proteomic profiling of low-input samples

Recognizing the benefits that could be attained from digestion in a droplet (*e.g.,* reduction in surface area contact and enhanced enzyme kinetics), we sought to develop an accessible platform for low-input proteomic sample preparation (Figure 1). We designed a microscope slide with a ∼20 µm coating of polytetrafluoroethylene (*i.e.,* Teflon) - a hydrophobic, chemically inert polymer – to contain aqueous droplets of ≤ 10 µL. The slides were procured from a commercial vendor with well spacing to match 96- well plates, enabling the use of standard multichannel pipettes. We designed and 3D-printed an adapter composed of polylactic acid that has the same footprint as a standard 96-well plate (available at https://3d.nih.gov/entries/3DPX-020483). This allowed us to easily transfer and position four DROPPS slides within plate-compatible flow cytometers, allowing parallel processing of 96 samples.

**Figure 1.**
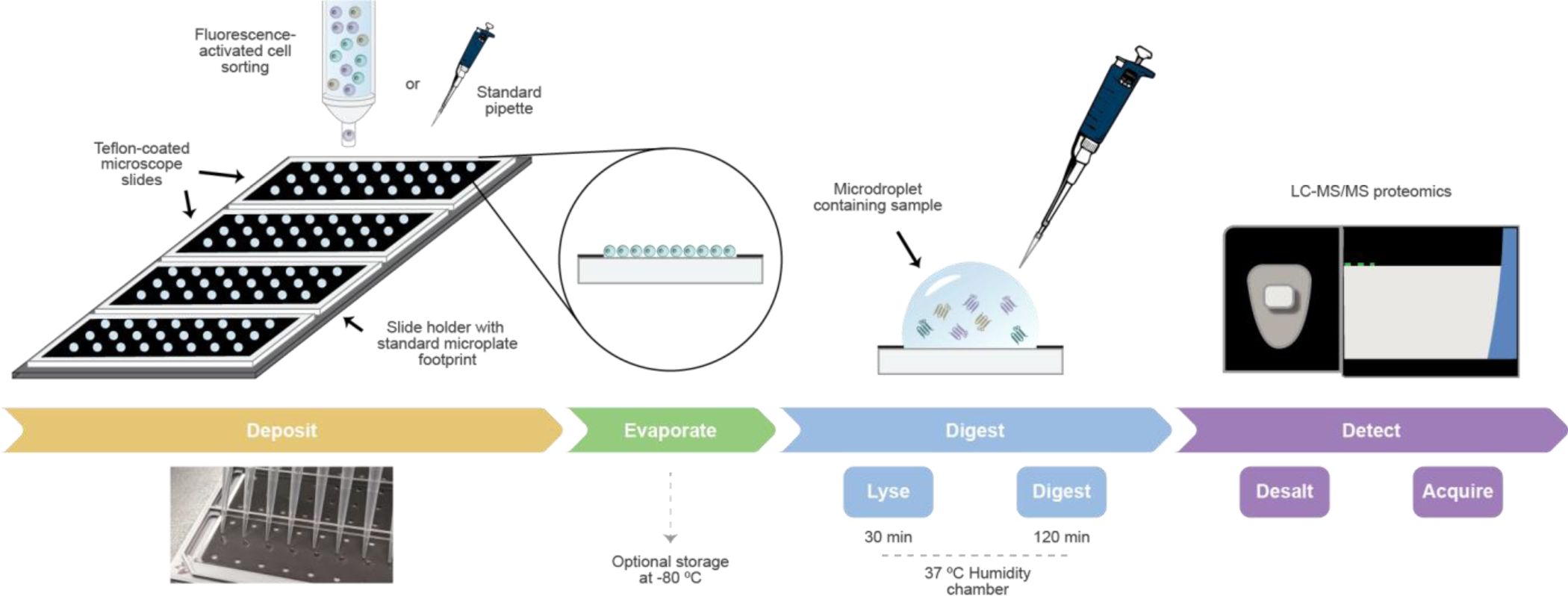
DROPPS workflow for low input protein digestion. Cells are deposited onto Teflon-coated slides by fluorescent-activated cell sorting or standard pipette. Cell transfer buffer is evaporated, and slides can be optionally stored at -80 °C. Cell lysis and protein digest is performed with MS-compatible reagents in a simple humidity chamber. Samples are transferred to standard sample vials or plates and analyzed by LC−MS for quantitative proteomic analysis.

To determine the range of sample input that can be processed by DROPPS with high reproducibility, we evaluated the performance of DROPPS over a range of starting material (10 – 2,500 FAC-sorted cells, *n* = 3) (Figure S1a) from MCF10A, a human mammary epithelial cell line. As expected, the number of input cells drastically affected the depth of proteomic coverage and variability in measurements (Extended Data Figure 1b-e). The proteomic depth obtained by DROPPS is a marked improvement (>50%) over other low-input proteomics methods.^19^ Protein intensity was linear across the 100-2,500 cell range supporting the utility of DROPPS for quantitative profiling rare populations of cells (plot shown for the luminal epithelial marker KRT18, median R^2^ of 0.97 Extended Data Figure 1f-g). We evaluated DROPPS reproducibility by having three operators perform independent tryptic digest of 500 MCF10A cells (*n* = 8) (Extended Data Figure 1h). To emulate typical workflows used for cellular biology, cells were counted by hemocytometer and deposited on the glass slide by pipette. The method was highly reproducible, with similar numbers of proteins and peptides detected in samples prepared by each operator (Extended Data Figure 1i-j). Technical variability from sample handling was low (median peptide coefficient of variation (CVs) ranging from 12.4% to 14.5% and protein CVs ranged from 8.3% to 9.9% per operator) (Extended Data Figure 1k-l). Overall, these results demonstrate that DROPPS and can be used to reproducibly generate quantitative proteomic profiles of 100 - 2,500 cells using common methods of sample handling.

### Biological differences of breast cancer cell lines are recapitulated by DROPPS profiles

We next evaluated the ability of DROPPS to repeatably detect biological differences among three human triple negative breast cancer (TNBC) cell lines (HCC1187, MDA-MB-157, and Hs-578-T) across three batches of 500 cell replicates (*n* = 7-8 per batch) (Figure 2a). To rigorously assess DROPPS performance across batches, every step of the workflow (*i*.*e*., cell culture, cell sorting, protein digestion, mass spectrometry) was completed on different days. Variance in quantitation was low across batches despite differences in the depth of proteomic coverage (Figure 2b-c, Extended Data Figure 2a-b). TNBC cell line protein profiles were highly similar within and between batches with median Spearman’s ρ of 0.984 and 0.970, respectively (Extended Data Figure 2c). Principal component analysis (PCA) distinctly separates the cell lines (Figure 2d). Differences between cell lines were larger than differences between batches for the majority (80%) of proteins (Figure 2e). These results show that most of the observed protein variance is due to biological differences between cell lines rather than experimental variance. Importantly, proteomic profile differences acquired by DROPPS were highly similar to proteomes acquired in two independent bulk proteomic datatsets^13,14^ (Spearman’s ρ of 0.62 and 0.64, both P < 2.2 x 10^-16^) (Extended Data Figure 2d-e). Furthermore, our data also successfully clustered breast cancer cell lines into molecular subtypes of TNBC^21^ indicating that the profiles generated by DROPPS accurately reflect previously described phenotypic differences (Figure 2f, Extended Data Figure 2f, Supporting Information Table 1). These data indicate our platform consistently and accurately recapitulates biological features of TNBC from 500 cells.

**Figure 2.**
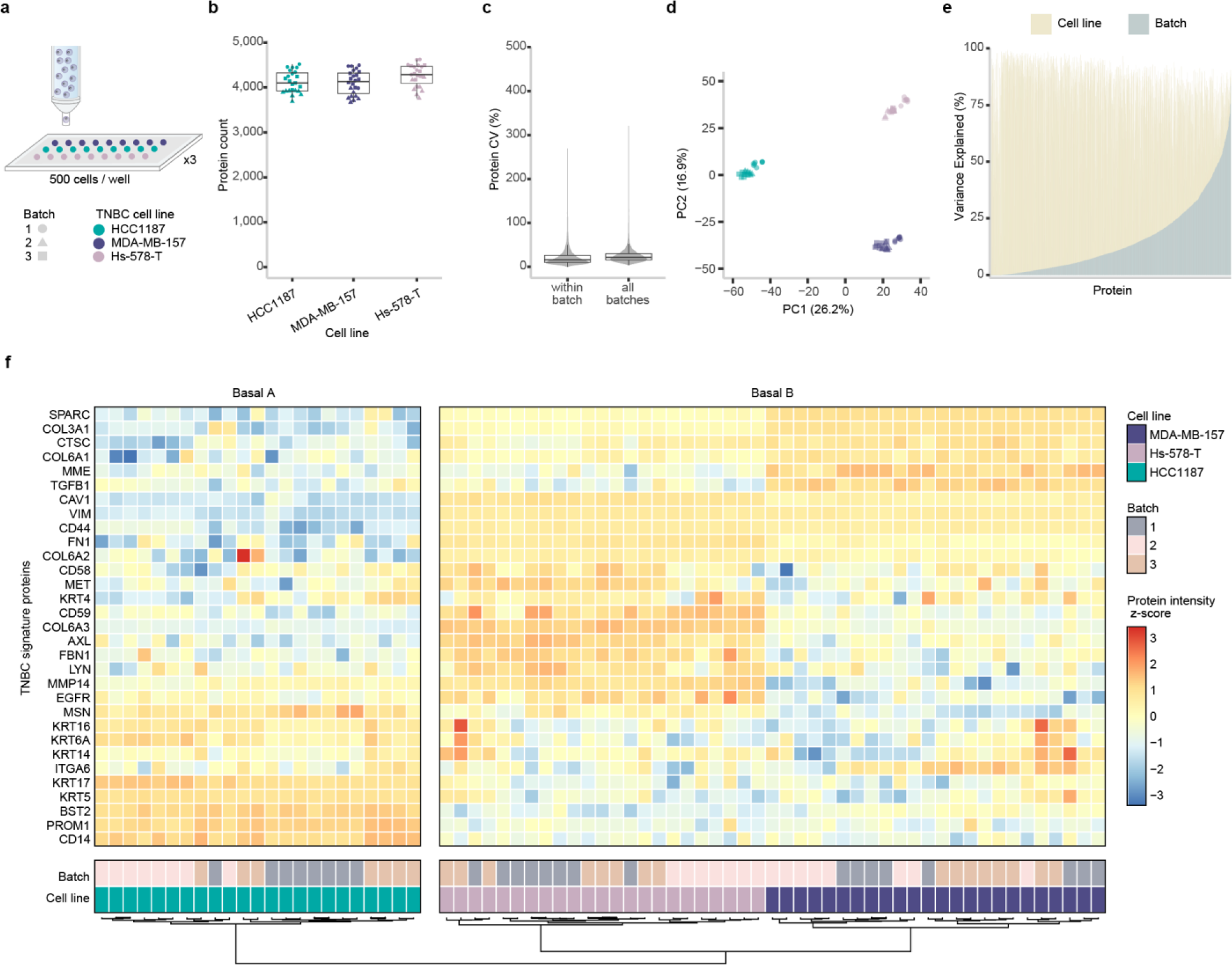
Evaluation of DROPPS for profiling TNBC cell lines differences across batches. (a) Workflow for testing inter-batch variation by DROPPS. Sorted cells (500 cells) from three TNBC cell lines were processed separately on different days. Each cell line had seven or eight sample replicates processed each day. (b) The protein counts of the individual runs. (c) Protein CVs for runs within a batch or for combining batches. (d) First two principal components of PCA. (e) The percentage of variance explained by cell line and batch from two-way ANOVA (*i.e.*, omega-squared). (f) z-scored log_2_(intensity) of TNBC signature proteins,^21^ using data acquired by DROPPS. Boxplots show the median, interquartile ranges, and 95% confidence interval estimate. TNBC: triple negative breast cancer

### Profiles of mammary epithelial compartments can be obtained from individual mice

The mammary epithelium is a ductal structure composed of an inner layer of luminal cells encompassed by basal cells – the cells from luminal and basal compartments have distinct molecular phenotypes which reflect their different functions.^20,22^ Here, by applying DROPPS we obtained improved depth of proteomic profiles for FACS purified luminal and basal compartments from individual mice than had previous achieved ^22^ by pooled samples (Figure 3a, Extended Data Figure 3a-d). Proteins uniquely detected in DROPPS were less abundant and detected in fewer mice highlighting heterogeneity among mice uncovered by the capacity to analyze ∼ 60-fold fewer cells than our previous study^22^ (Extended Data Figure 3e-f). As expected, differential expression analysis revealed pronounced differences between luminal and basal cells that were highly correlated to our previous study^22^ (Figure 3b, Extended Data Figure 3g). The cell surface proteins used to distinguish luminal from basal cells^23–25^, ITGA6 (CD49f) and EPCAM, were among the most significantly different along with other described lineage markers^22^ (*e.g.*, KRT5/14 and KRT8/18/19) (Figure 3b). These results show that DROPPS enables the investigation of mammary epithelial subpopulations (of at least 2,000 cells) from individual mice - a significant advance enabling the investigation of proteomic heterogeneity within and between samples.

**Figure 3.**
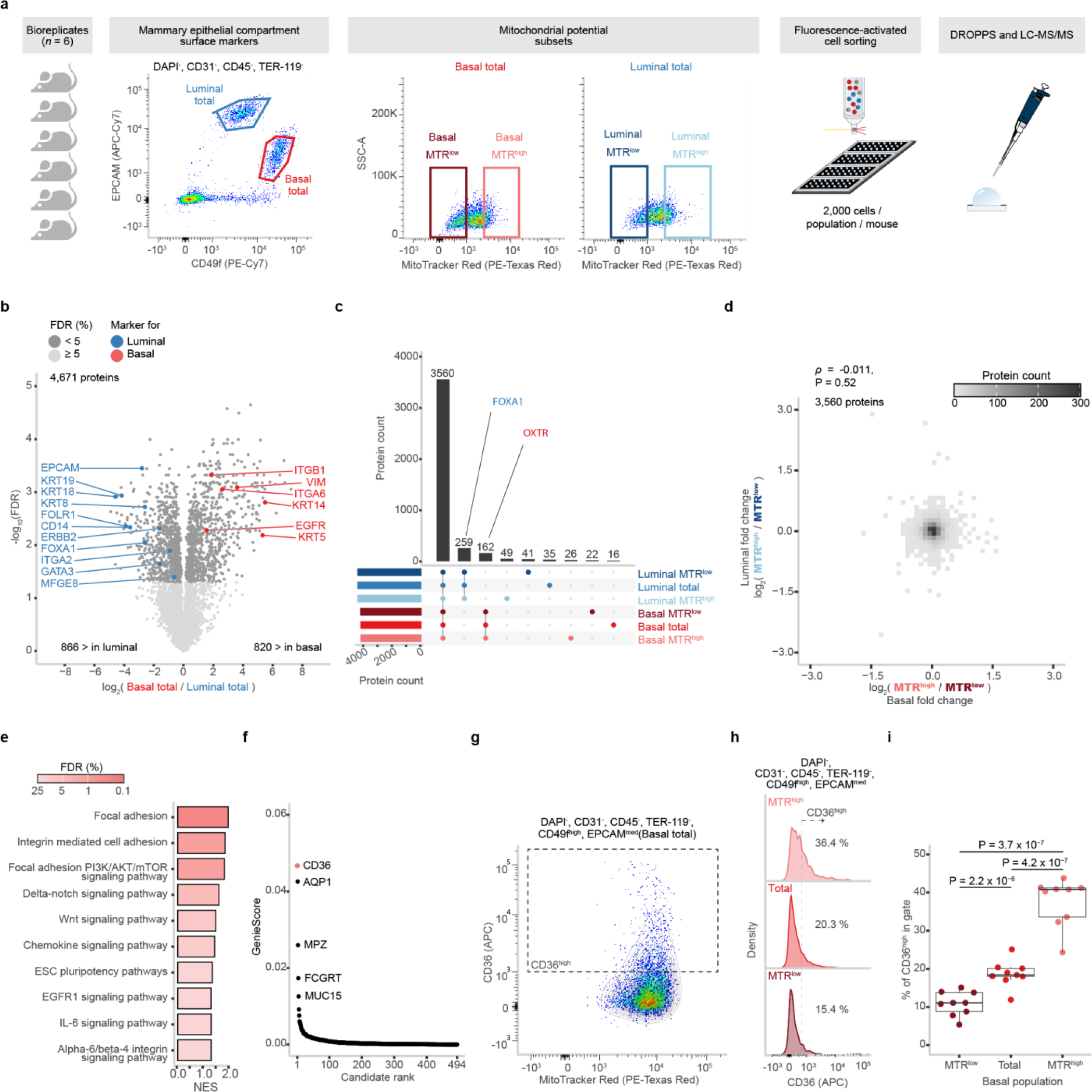
Proteomic profiling of primary mammary epithelial cells sorted by mitochondrial potential. (a) Schematic of experimental design and FACS gating scheme used to sort subsets of mammary epithelial cells of basal and luminal compartments based on their mitochondrial potential as marked by MTR dye (*n* = 6 each) that were subsequently processed using DROPPS. (b) Differences in log_2_(intensities) of proteins from the total basal and total luminal populations. Points are colored by significance and known compartment markers are labeled (c) Overlap in proteins detected between select subsets of epithelial cells. (d) Two-dimensional bin plot depicting the distribution of log_2_ fold-changes between MTR^high^ and MTR^low^ for the basal and luminal compartments. (e) Select GSEA enriched pathways in the MTR^high^ population compared to the total population for the basal and luminal compartments. (f) Rank plot depicting the distribution of GenieScores for proteins with Surface prediction consensus > 0 calculated using SurfaceGenie^29^ on mean log_2_(intensities). (g) Representative scatterplot depicting the relationship between CD36 and MTR fluorescent intensity. (h) Representative histograms depicting the distribution of CD36 flow cytometry signal in MTR^low^, total, and MTR^high^ basal populations. (i) The percentage of CD36^high^ cells in different MTR-fractions of basal cells shown with paired t-test p- values (*n* = 8). Boxplots show the median, interquartile ranges, and 95% confidence interval estimate. MTR: MitoTracker Red.

### CD36 identified as a marker of basal mitochondrial potential

The ability to monitor and target oncogenic precursors could improve clinical outcomes for breast cancer patients.^8^ As stem/progenitor cells give rise to the deadliest breast cancer subtypes,^5–7^ it is of particular interest to identify biomarkers and molecular vulnerabilities of these cell types. We previously identified a positive correlation between mitochondrial membrane potential (ΔΨ_M_) - measured by MitoTracker Red (MTR) - and clonogenicity within mammary epithelial compartments.^20^ Luminal and basal compartments both had significantly diminished clonogenicity within MTR^low^ fractions. MTR^high^ fractions demonstrated a modest increase of clonogenicity in the luminal compartment and a significant increase in the basal compartment. As clonogenicity is an established trait of stem/progenitor cells, we applied DROPPS to FACS purified MTR^high^, MTR^low^, and total cell fractions from basal and luminal compartments to interrogate how ΔΨ_M_ heterogeneity relates to clonogenicity within the mouse mammary epithelium (2,000 cells per fraction, *n* = 6) (Figure 3a, Extended Data Figure 3a). Overall, we detected a total of 5,576 proteins (a mean of 4,423 proteins per sample) (Extended Data Figure 3b, 3d). As expected, we revealed a substantial overlap in the proteomes with some known compartment markers demonstrating compartment-specific detection (*e.g*., FOXA1^23,26^, OXTR^27,28^) (Figure 3c, Supporting Information Table 1). Unsurprisingly, epithelial compartment was the largest source of sample variance (Extended Data Figure 3h).

The proteins correlated with ΔΨ_M_ were completely independent between the luminal and basal cells (Spearman’s ρ of -0.011, P = 0.52) suggesting compartment-specific pathways associated with MTR^high^ and MTR^low^ populations (Figure 3d). We performed differential expression analysis to interrogate which pathways are associated with MTR^high^ in each compartment. Luminal-MTR^high^ cells are enriched in oxidative phosphorylation - the cellular process that generates the proton gradient component of ΔΨ_M_ (Supporting Information Table 2). In contrast, basal-MTR^high^ cells are enriched in various proliferation, survival, adhesion-sensitive growth, and pluripotency pathways (Supporting Information Table 3, select terms in Figure 3e). These results align with the previously described^20^ differences, namely, a larger effect on clonogenicity in the basal compartment for MTR^high^ populations. The accessibility of surface proteins makes them attractive markers and drug targets. As basal MTR^high^ cells are highly clonogenic and have progenitor associated pathways, we used SurfaceGenie^29^ to rank potential cell surface markers revealing CD36 as a high-priority, differentially expressed candidate (Figure 3f, Extended Data Figure 3i). We validated that CD36 expression was positively correlated with ΔΨ_M_ by live cell flow cytometry (Figure 3g) and observed significantly different frequencies of CD36^high^ cells between MTR^low^, total, and MTR^high^ subsets (Figure 3h-i). These results clearly depict a relationship between ΔΨ_M_ and CD36 in basal epithelial cells. Overall, these results demonstrate how DROPPS can be applied to generate insights into mammary biology and to identify cell surface markers for desired epithelial subpopulations.

### Basal progenitors are enriched by CD36

We next investigated whether CD36 was associated with enhanced clonogenicity, or solely ΔΨ_M_, by comparing CD36^high^ cells to the total basal population using a colony forming assay (Figure 4a). CD36^high^ cells had enhanced clonogenicity shown by significantly more colonies (paired t-test, P = 0.039) and larger colonies (paired t-test, P = 0.048) (Figure 4a-c, Extended Data Figure 4a). We then analyzed CD36^high^ basal cells by DROPPS to determine which pathways were associated with the enhanced clonogenicity detecting a mean of 4,187 proteins per sample (2,000 cells per mouse, *n* = 8) (Extended Data Figure 4b). We detected 49 proteins significantly enriched in CD36^high^ compared to the total basal population – including CD36 (Figure 4d). CD36 expression was associated with enriched proliferation (*e.g.*, growth and development, cell division, stem cell proliferation), various signaling pathways (*e.g.*, Notch, MAPK, Rho, NF-κB), and metabolism (*e.g.*, organic hydroxy compound transport, monocarboxylic acid transport, and lipid homeostasis) – all potentially contributing to the enhanced clonogenicity of the CD36^high^ population (Supporting Information Table 2, clustered terms in Figure 4e). A major function of CD36 is the uptake of long chain fatty acid which has been tied to survival, metabolic rewiring, and epithelial-mesenchymal transition in breast cancer.^30–32^ We investigate whether CD36 was associated with increased fatty acid accumulation in basal epithelial cells using imaging flow cytometry with a fluorescent long-chain fatty acid analog, BODIPY-dodecanoic acid (Figure 4g-I, Extended Data Figure 4d). Our analyses revealed that CD36 expression in basal cells was associated with significantly increased BODIPY signal indicating a potential mechanism by which CD36 contributes to increased ΔΨ_M_ and enhanced progenitor capacity (Figure 4j-k). Taken together, these discoveries exhibit how DROPPS can provide insights to the underlying biology of rare cell populations.

**Figure 4.**
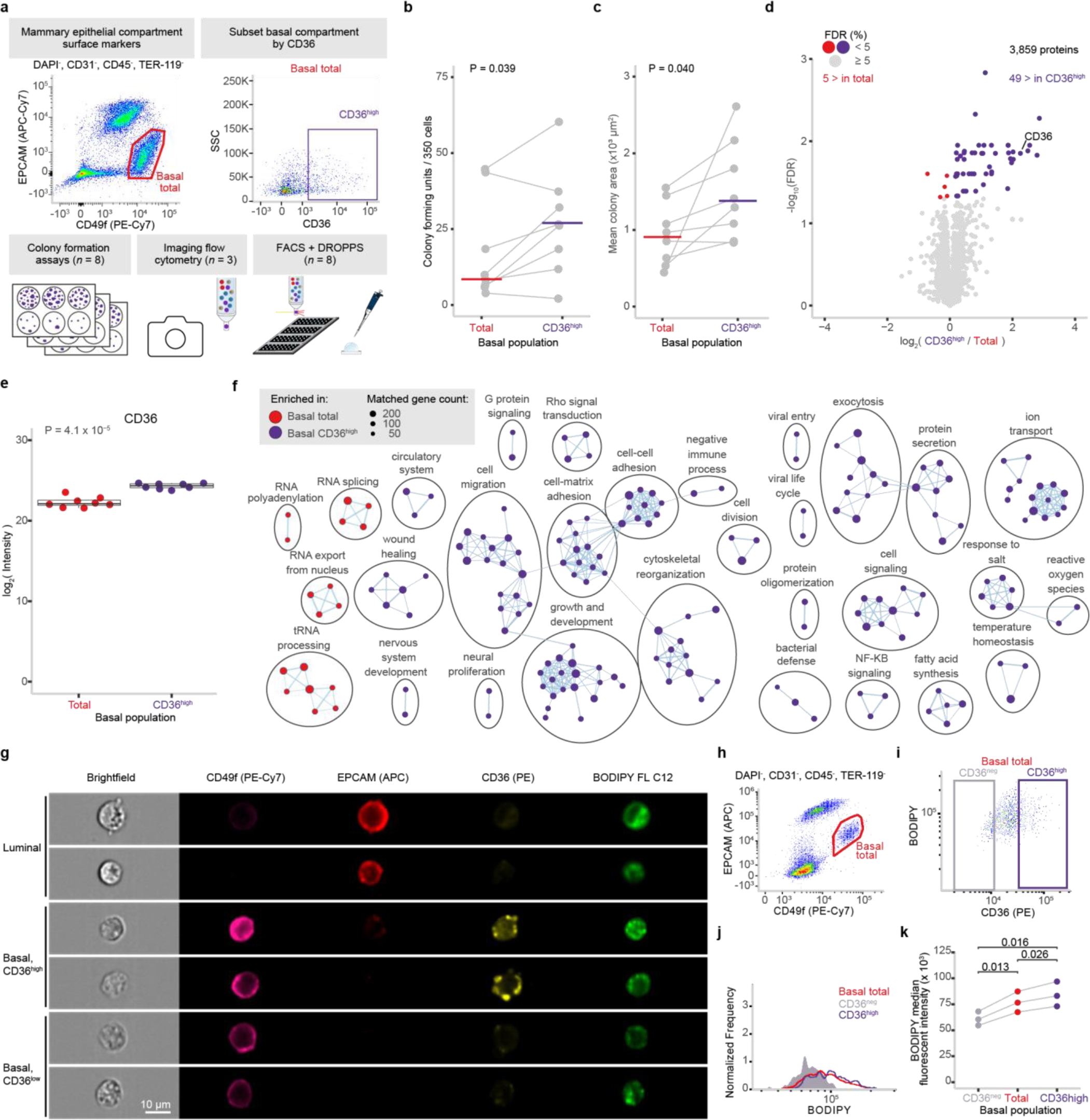
Investigating the phenotype of CD36^high^ basal mammary epithelial cells. (a) Schematic of experimental design and FACS gating scheme used to sort subsets of mammary epithelial cells of basal and luminal compartments based on CD36 expression that were subsequently used for CFC, processed using DROPPS, or analyzed by imaging flow cytometry. (b,c) Line plots depicting the (b) number and (c) size of the colonies formed by basal cells sorted by CD36 expression shown with median value and paired t-test p-values. (d) Differential protein expression results from total basal and CD36^high^ basal populations. Points are colored by significance and directionality. (e) The log_2_(intensity) of CD36 from individual runs shown with the paired t-test p-value (*n* = 8). (f) Cytoscape enrichment map for significantly enriched GO Biological Processes between total basal and CD36^high^ basal populations population. (g) Representative images from imaging cytometry highlighting the surface markers used to distinguish epithelial compartments along with CD36 and the fluorescent fatty acid analog BODIPY. (h) Scatterplot depicting the gating scheme used to identify basal epithelial cells. (i) Scatterplot depicting the relationship between BODIPY and CD36 intensities along with CD36^neg^ and CD36^high^ gates. (j) Representative histogram depicting the BODIPY intensity distributions among CD36^neg^, total, and CD36^high^ basal populations (k) Dot plot depicting the median BODIPY fluorescent intensity in CD36^neg^, total, and CD36^high^ basal populations along with paired t-test p-values. CFC: colony forming capacity.

## Discussion

Unraveling the mechanisms underlying development, proper function, and disease will require understanding the dynamic changes and interactions of myriad cell types including those that are rare. Here, we developed and validated DROPPS, a proteomic workflow designed to democratize the capacity to profile rare cell populations without a compromise of data quality. DROPPS generates high fidelity proteomic profiles from samples comprising as low as 100 cells and is amenable to facile integration with FACS. DROPPS performance is robust across operators and sample processing batches, and it is easy to implement - relying exclusively on common or affordable laboratory equipment. Provided access to liquid handling systems, automation of DROPPS should be practical as it involves very limited sample handling and straightforward transfer steps. The capacity to rapidly generate protein profiles of rare cell populations enables experimental designs which were previously inaccessible or impractical.

There remains discrepancy regarding the exact differentiation hierarchy of mammary epithelial cells and how to identify and enrich for populations of different functions along the lineage trajectory. Here, we applied DROPPS to examine the proteomic correlates of a previously described^20^ relationship between ΔΨ_M_ and clonogenic potential in lineage-defined mouse mammary epithelial cells. Specifically, basal- MTR^high^ cells demonstrated a significant increase in clonogenicity compared to the total basal population. MTR^low^ cells in both basal and luminal compartments had significantly diminished clonogenicity. DROPPS revealed that differences in ΔΨ_M_ were reflected in the proteome in a compartment-specific manner. Basal-MTR^high^ cells were enriched in pathways associated with proliferation and adhesion-dependent signaling. We uncovered CD36 as a candidate marker for basal cells with elevated ΔΨ_M_ and enhanced clonogenicity. This work contributes to our understanding of the relationship between molecular phenotype and clonogenicity in the mammary basal epithelium and provides a marker, CD36, by which primary cells with progenitor capacity can be enriched and studied.

CD36 is a multifunction cell surface glycoprotein that acts as a receptor for a broad range of ligands including proteins and lipids. In this study, we found CD36 was associated with clonogenicity, increased fatty acid abundance, and enrichment of myriad growth, signaling, and metabolic pathways in primary basal epithelial cells. The putative association between increased fatty acid uptake and progenitor capacity aligns with our single cell RNA-Sequencing analysis of human mammary basal epithelial cells, in which higher oxidative phosphorylation was associated with a less differentiated cell state.^20^ In another study that characterized features of mouse mammary stem cells, CD36 is among the proteins associated with the mammary stem cell signature.^33^ The expression of CD36 is associated with increased free fatty acid uptake and metabolic rewiring in a variety of biological systems including hepatocellular carcinoma, hematopoietic stem cells, and breast cancer.^30,34,35^ CD36 has also been implicated in enhancing proliferation, migration, and epithelial-to-mesenchymal transition in multiple types of cancer.^32,36,37^ Our studies identify CD36 a marker for progenitor capacity in basal mammary epithelium that may function to metabolically reprogram cells towards a less differentiated and proliferative state.

In summary, we present an accessible proteomic platform, DROPPS, and demonstrate its utility to uncover biological insights of rare epithelial cell populations. DROPPS affords the capacity to assess mouse-to-mouse variability, differences between subsets of cells within each mouse, and rare populations of cells thus enabling proteomic insights into the functional heterogeneity that were precluded by previous sample requirements. DROPPS has widespread applicability to various sample-limited systems and is well positioned to transform our ability to map the proteomic landscapes of rare cell populations.

## Methods

### Experimental model details

Cell lines: All cell lines used in this study were purchased from ATCC, were authenticated using short tandem repeat (STR) DNA profiling identity at TCAG Facilities (Sick Kids Hospital, Toronto), and were tested for mycoplasma contamination using the ATCC universal mycoplasma detection kit according to the instructions of the manufacturer. Growth media and supplements were purchased from Gibco unless otherwise specified. Hs 578T (ATCC® HTB-126™) and MDA-MB-157 cells were cultured in DMEM media supplemented with 10% fetal bovine serum (FBS) and penicillin-streptomycin (100 U/mL penicillin, 100 μg/mL streptomycin, Gibco). HCC1187 (ATCC® CRL-2322™) cells were cultured in RPMI media supplemented with 10% FBS and penicillin-streptomycin (100 U/mL penicillin, 100 μg/mL streptomycin). MCF10A (ATCC® CRL-2322™) cells were grown in DMEM:F12 media supplemented with 5% horse serum, 20 ng/mL epidermal growth factor (EGF), 10 μg/mL insulin, 500 ng/mL hydrocortisone, 100 ng/mL cholera toxin, and penicillin-streptomycin (100 U/mL penicillin, 100 μg/mL streptomycin). All cell lines were incubated at 37°C, 5% CO_2_.

Mouse experiments: All experiments were performed using 8- to 12-week-old virgin female FVB wild-type mice (The Jackson Laboratory or Charles River). Mice were ovariectomized bilaterally, then allowed 10- 14 days to recover. A slow-release 0.14 mg 17-β oestradiol plus 14 mg progesterone pellet (Innovative Research of America) was then placed subcutaneously near the thoracic mammary gland for 12-14 days. This was done to obtain large quantities of viable mammary stem/progenitor cells for subsequent analysis, as previously reported.^22,38^ The light cycle in the rooms was set to 12 h on and 12 h off with a 30-min transition. Temperature was set to 21–23 °C and humidity at 30–60%. All animal procedures were in compliance with ethical guidelines established by the Canadian Council for Animal Care under protocols approved by the Animal Care Committee of the Ontario Cancer Institute.

### Mouse mammary single-cell suspensions

Mammary glands were collected and manually minced with scissors for 2 min, and then enzymatically dissociated using 750 U/mL collagenase and 250 U/mL hyaluronidase (STEMCELL Technologies, 07912) and diluted in DMEM:F12 for 1.5 h. Samples were vortexed after 1 h and 1.5 h. Red blood cells were lysed using ammonium chloride (STEMCELL Technologies, 07850). Cells were then triturated in prewarmed (37 °C) trypsin-EDTA (0.25%; STEMCELL Technologies, 07901) with a 1 mL pipette for 2 min. Next, they were washed in HBSS without calcium or magnesium plus 2% FBS and centrifuged at 350 x *g*. Finally, cells were further dissociated in Dispase 5 U/mL (STEMCELL Technologies, 07913) plus 50 µg/mL DNase I (Sigma, D4513) for 2 min, washed in HBBS + 2% FBS and filtered using a 40 µm cell strainer to obtain single cells.

### Cell line single-cell suspensions

Single cell suspensions were generated by washing cells with PBS followed by 3-5 min incubation in TryplE. Digestion was quenched with equal volume of growth media and collected by centrifugation at 300 x *g*. Cells were washed twice with PBS before incubation with DAPI. After 10 min incubation, cells were collected by centrifugation at 300 x *g* and filtered through a 35 μm cell strainer.

### Fluorescent activated cell sorting

Cell lines: Dead cells were excluded using DAPI following doublet exclusion.

Mouse experiments: Staining for mitochondrial activity (250 nM MTR CM-H2Xros; ThermoFisher, M7513) was performed before cell-surface-marker staining protocol by incubating cells at 37 °C for 30 min following the manufacturer’s protocols. For FACS staining, APC-Cy7 rat anti-mouse CD326 clone G8.8 (EpCAM; BioLegend, 118217; 1:200 dilution), PE-Cy7 rat anti-human CD49f clone GoH3 (BioLegend, 313622; 1:100 dilution), eFluor™ 450 rat anti-mouse CD31 clone 390 (ThermoFisher, 48-0311-82; 1:200 dilution), eFluor™ 450 rat anti-mouse CD45 clone 30-F11 (ThermoFisher, 48-0451-82; 1:800 dilution), eFluor™ 450 rat anti-mouse TER-119 clone TER-119 (ThermoFisher, 48-5921-82; 1:100 dilution), APC Mouse Anti-Mouse CD36 clone CRF D-2712 (BD Biosciences, 562744; 1:20 dilution) were used. Dead cells were excluded following doublet exclusion using DAPI. Lineage-positive cells were defined as Ter119^+^CD31^+^CD45^+^. Mouse mammary cell subpopulations were defined as: total basal (Lin^−^EpCAM^lo-med^CD49f^hi^), total luminal (Lin^−^EpCAM^hi^CD49f^lo^) as previously described.^23–25^ MTR^high^ and MTR^low^ mitochondrial activity populations were defined as top 10 % and bottom 30 % of MTR signal, respectively, and applied after gating for total luminal and basal populations. Cell sorting was performed on a BD FACS ARIA Fusion with FACSDiva (v.8.0.1 and v.6.1.3). Flow data analysis was performed using FlowJo (v.10.8.1).

### Imaging flow cytometry

Incubation of samples in 1 µM BODIPY™ FL C12 (D3822, ThermoFisher) was performed for 30 min at 37 °C for quantification of fatty acid prior to incubation with cell surface antibodies. Cell surface marker staining was performed using the same antibodies against CD49f, CD31, CD45, and TER-119 as in fluorescent activated cell sorting experiments. In addition, APC rat anti-mouse CD326 clone G8.8 (EpCAM; BioLegend, 118214; 1:200 dilution) and PE Mouse Anti-Mouse CD36 clone CRF D-2712 (BD Biosciences, 562702; 1:20 dilution) were used. CD36^high^ was annotated as the top ∼20% CD36 expressing cells and CD36^neg^ was set using fluorescent-minus-one control. Raw imaging flow cytometry files were captured on a Cytek ImageStream AMNIS MKII using INSPIRE software and data analysis performed using IDEAS 6.3 software.

### Mouse colony-forming cell assay

In total, 350 cells of the specified FACS-purified population were seeded together with 20,000 irradiated NIH 3T3 mouse fibroblasts per well in six-well plate format. Cells were cultured for 7 d at 5% O_2_ in EpiCult- B mouse medium (STEMCELL Technologies, 05610) supplemented with 5% FBS, 10 ng/ml human epidermal growth factor (STEMCELL Technologies, 78006), 20 ng/mL basic fibroblast growth factor (STEMCELL Technologies, 78003), 4 μg/mL heparin (STEMCELL Technologies, 07980) and 5 μM ROCK inhibitor (Millipore). Colonies were stained with Wright-Giesma and plates were scanned using the Biotek Cytation 5 Imaging Multimodal Reader (Agilent). From scanned images, colonies were counted and sized using Biotek Gen5 software (v3.11).

### Proteomic sample preparation

Cells were deposited into wells of Teflon-coated slides (Tekdon, template available upon request) by pipette or directly by FACS. The solution accompanying the cells was allowed to evaporate at room temperature in a biosafety cabinet and then slides were stored at -80 °C until further processing. For lysis and protein reduction, 2.5 μL of buffer consisting of 30% (v/v) Invitrosol (ThermoFisher, MS10007), 15% (v/v) acetonitrile, 0.06% (w/v) n-dodecyl-β-D-maltoside (Sigma, 850520P), 5 mM Tris(2- carboxyethyl)phosphine, and 100 mM ammonium bicarbonate in HPLC grade water. Slides were placed in a 37 °C humidity chamber and allowed to incubate for 30 min. For alkylation and protein digestion, 1 μL of buffer containing 35 mM iodoacetamide and Trypsin/Lys-C (Promega, V5072) was added to each sample. Samples were allowed to digest for 2 h in a 37 °C humidity chamber. Digestion was quenched by bringing samples to 0.1% (v/v) formic acid with 8 μL of 0.14% (v/v) formic acid in 37 °C HPLC grade water and then samples were transferred to 96-well plate. The wells were washed with an additional 8 μL of 0.1% (v/v) formic acid and then the plate was transferred to EASY-nLC™ 1000 System chilled to 7 °C.

### Mass spectrometry data acquisition

LC-MS/MS analysis was performed on an Orbitrap Fusion MS (ThermoFisher) coupled to EASY-nLC™ 1000 System (ThermoFisher). Peptides were washed on pre-column (Acclaim™ PepMap™ 100 C18, ThermoFisher) with 60 μL of mobile phase A (0.1% FA in HPLC grade water) at 3 μL/min separated using a 50 cm EASY-Spray column (ES803, ThermoFisher) ramping mobile phase B (0.1% FA in HPLC grade acetonitrile) from 0% to 5% in 2 min, 5% to 27% in 160 min, 27% to 60% in 40 min interfaced online using an EASY-Spray™ source (ThermoFisher). The Orbitrap Fusion MS was operated in data dependent acquisition mode using a 2.5 s cycle at a full MS resolution of 240,000 with a full scan range of 350-1550 *m/z* with RF Lens at 60%, full MS AGC at 200%, and maximum inject time at 40 ms. MS/MS scans were recorded in the ion trap with 1.2 Th isolation window, 100 ms maximum injection time, with a scan range of 200-1400 *m/z* using Normal scan rate. Ions for MS/MS were selected using monoisotopic peak detection, intensity threshold of 1,000, positive charge states of 2-5, 40 s dynamic exclusion, and then fragmented using HCD with 31% NCE.

### Mass spectrometry raw data analysis

Raw files were analyzed using FragPipe (v.20.0) using MSFragger^39,40^ (v.3.8) to search against a human (Uniprot, 43,392 sequences, accessed 2023-02-08) or mouse (Uniprot, 25,474 sequences, accessed 2021-06-03) proteomes – canonical plus isoforms. Default settings for LFQ workflow^41,42^ were used using IonQuant^43^ v.1.9.8 and Philosopher^44^ v.5.0.0 with the following modifications: Precursor and fragment mass tolerance were specified at -50 to 50 ppm and 0.15 Da, respectively; parameter optimization was disabled; Pyro-Glu or loss of ammonia at peptide N-terminal was included as a variable modification; MaxLFQ min ions was set to 1; MBR RT tolerance was set to 2 min, and MBR top runs was set to 100. The cell titration experiment was searched without MaxLFQ and without normalization.

### Mass spectrometry statistical analysis

All analysis was performed using R programming language (v.4.2.2) with Tidyverse pacakge (tidyverse_1.3.2) unless otherwise specified. All correlation estimates and p-values were calculated using the “cor.test” function. For all experiments except the titration experiment, the “MaxLFQ Intensity” columns were extracted from the “combined_protein.tsv” and “combined_peptide.tsv” output files from FragPipe (Supporting Information Table 3). For the titration experiment the “Intensity” columns were used. Proteins were filtered out if the Uniprot Accession ID matched one from the MaxQuant contaminants list unless they belonged to a set of IDs being used for comparison (*e.g.*, TNBC Basal subtype markers or epithelial compartment markers, Supporting Information Table 1). Coefficients of variation were calculated on intensity measurements with three or more replicates without imputation. Run-to-run correlations were calculated using Spearman’s correlation on log_2_-transformed intensities without imputation. Principal component analysis was performed on log_2_-transformed intensities imputing missing values to 0.

TNBC cell line analysis: First proteins were filtered for presence in > 50% of replicates for at least one cell line and subsequently imputed with lower tail imputation (downshift of 1.8 s.d. and width of 0.5 s.d.) ^45^. TNBC basal markers^21^ heatmap generation was clustered using Pretty Heatmap package (pheatmap_1.0.12) using correlation as a distance metric for proteins, Euclidean distance for samples, and Ward D2 linkage The GSVA^46^ package (GSVA_1.46.0) was used to calculated similarity between cell lines and TNBC Basal subtypes. Omega-squared values were calculated fitting log_2_-transformed intensities to a two-way ANOVA with batch and cell line as independent factors. Differential expression was calculated using the Estimated Marginal Means package (emmeans_1.8.4-1) to calculate group means from a linear model fit with batch as cell line as factors with an interaction term. Mean fold-changes were calculated from supplementary files from previous bulk proteomic studies^13,14^ and plotted against mean fold changes from this study.

Primary total epithelial compartments: First proteins were filtered for presence in > 50% of replicates for at least one epithelial compartment and subsequently imputed with lower tail imputation^45^. Differential expression was calculated using Estimated Marginal Means package using a linear mixed model with compartment as a fixed effect and allowing for a different intercept per mouse. Genes in volcano plot were annotated with canonical compartment markers as previously described.^22^ Data from Casey *et al.* were accessed from Table S5 and fold changes were calculated using the “LFQ or adj iBAQ” values.

Primary epithelial MTR subsets: Data for basal and luminal compartments were analyzed independently. First proteins were filtered for presence in > 50% of replicates for total, MTR^high^ and MTR^low^ subsets and subsequently imputed with random forest algorithm using the MissForest package (missForest_1.5). Significant differences were detected using repeated measures ANOVA with mitochondrial subset as variable within mouse as a grouping variable. Fold changes were estimated using Estimated Marginal Means package using a linear mixed model (lme4_1.1-31) with mitochondrial subset as a fixed effect and allowing for a different intercept per mouse. GSEA was applied using a preranked list of Gene Symbols sorted based on estimated fold changes against the Mouse Wikipathways gene sets with minimum and maximum sizes of 25 and 200, respectively. GenieScore was calculated as described^29^ using mitochondrial subset means as input values.

Basal CD36 population: First proteins were filtered for presence in > 50% of replicates for total and CD36^high^ subsequently imputed using the MissForest package. Fold changes and significance were estimated using Estimated Marginal Means package using a linear mixed model with CD36 subset as a fixed effect and allowing for a different intercept per mouse. GSEA was applied using a preranked list of Gene Symbols sorted based on estimated fold changes against the Mouse Gene Ontology Biological Processes gene sets with minimum and maximum sizes of 25 and 200, respectively. Outputs from GSEA were used as inputs to Cytoscape (v.3.9.1). GSEA gene sets were visualized in EnrichmentMap (v.3.3.4) and AutoAnnotate (v.1.3.5).

### Data visualization

Unless otherwise specified, plots were generated using R programming language (v.4.2.2) with Tidyverse pacakge (tidyverse_1.3.2) with ggthemes_4.2.4, ggpubr_0.5.0, ggplot2_3.4.0, and ggbeeswarm_0.6.0 packages.

## Acknowledgements

Work in the Kislinger and Khokha labs was supported by operating grants from the Canadian Institutes of Health Research, the Terry Fox Research Institute PPG, Canadian Cancer Society, the Canadian Research Chair program. A.K. was supported by the Ontario Graduate Scholarship and Ontario Student Opportunity Trust Fund Awards. CM was supported by a Banting Postdoctoral Fellowship. Flow cytometry was performed using instrumentation in the Princess Margaret Flow facility. We are especially thankful to Frances Tong, Aleks Spurmanis, and Arian Khandani for feedback on slide and plate design and assistance with sample collection. Special thanks to Patrik Lanzhotsky for assistance with 3D printing.

## Author Contributions

M.W., A.K., P.T., C.W.M., M.G., B.Z., S.K., and P.W. designed experiments, performed experiments, and contributed to data acquisition. M.W. and S.K. acquired the mass spectrometry data. M.W., A.K., P.T., C.W.M., and B.Z. analyzed the data. M.W. and A.K. generated figures. R.K. and T.K. supervised the study. M.W., A.K. and M.G. wrote the manuscript, which all other authors edited and approved.

## Competing interests

Authors have no competing interests to declare.

## Additional information

Supplementary Information is available for this paper. Correspondence and requests for materials should be directed to and will be fulfilled by lead contacts, Rama Khokha. Thomas Kislinger.

## Data availability

All mass spectrometry raw files and processed result files acquired in this study are publicly available from UCSD’s MassIVE database (ftp://massive.ucsd.edu) under dataset identifier MSV000092829. Processed proteomics data are available in this paper’s Supplementary Table 3. Any additional information required to reanalyze the data reported in this paper is available from the lead contact upon request.

## Supplementary Information

Supplementary Table 1: Protein and Gene Lists, related to all Figures

Supplementary Table 2: GSEA Results, related to Figure 3 and 4

Supplementary Table 3: Protein and Peptide Tables, related all Figures

Supplementary Table 4: Results from statistical analyses, related to Figures 3 and 4

---

**Figure S1.**
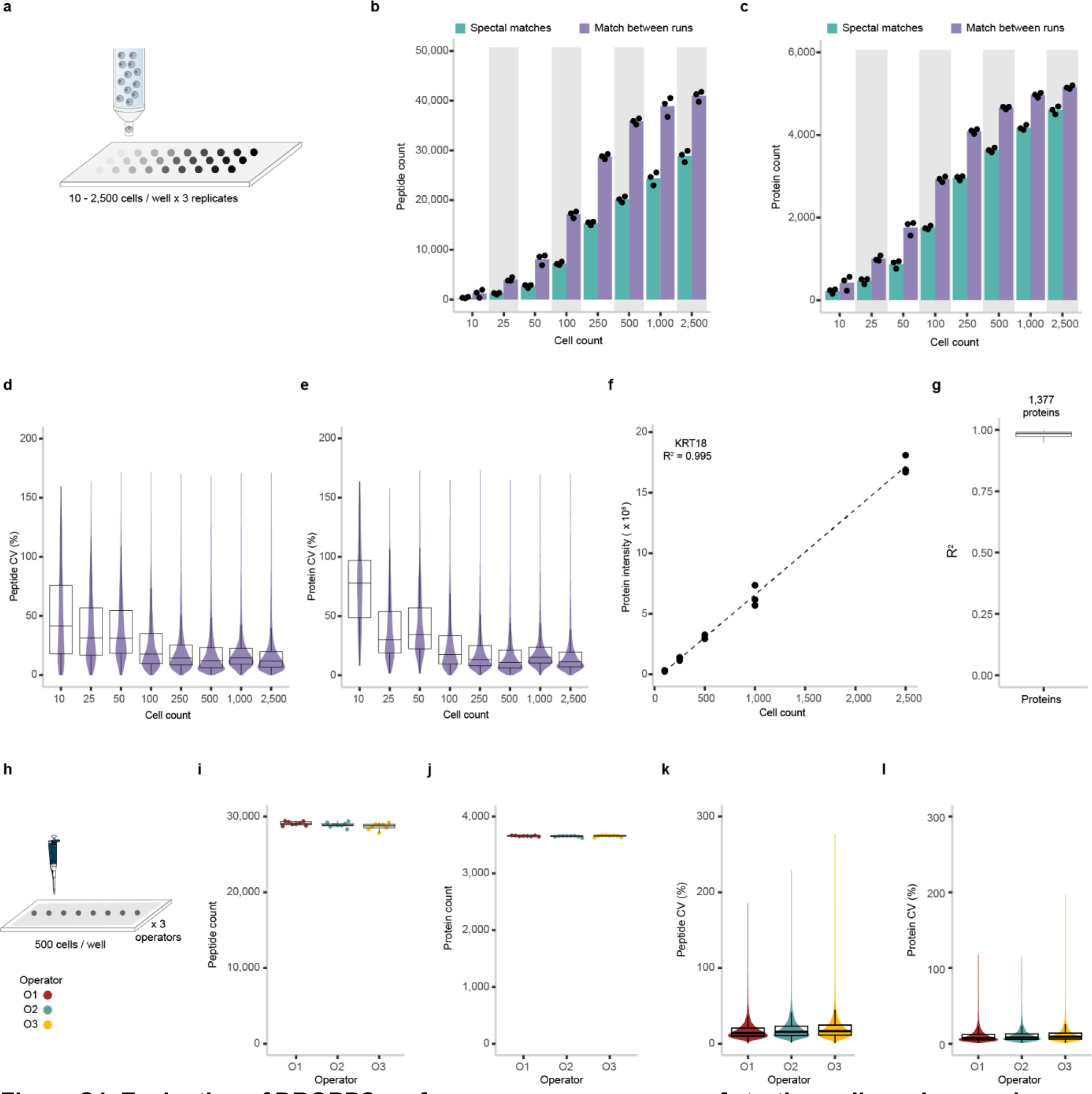
Evaluation of DROPPS performance across a range of starting cell numbers and among operators. (a) Cell number titration workflow where 10 – 2,500 MCF10A cells were deposited per well (*n* = 3 each) by fluorescent-activated cell sorting and subsequently processed using DROPPS. (b,c) The (b) peptide counts and (c) protein counts of the individual runs with and without “match between runs” where columns represent the mean values. (d,e) Distribution of intensity CVs at the (d) peptide and (e) protein level. (f) KRT18 protein intensity as a function of input cell number. (g) Distribution of R^2^ calculated from modeling the protein intensity as a function of cell number. (h) Schematic of operator experimental design where 500 MCF10A cells were deposited per well (*n* = 8 each) by three operators and subsequently processed using Dropps. (i,j) The (i) peptide counts and (j) protein counts of the individual samples prepared by each operator. (k,l) Distribution of (k) peptide intensity or (l) protein intensity CVs. Boxplots show the median, interquartile ranges with 95% confidence interval estimate.

**Figure S2.**
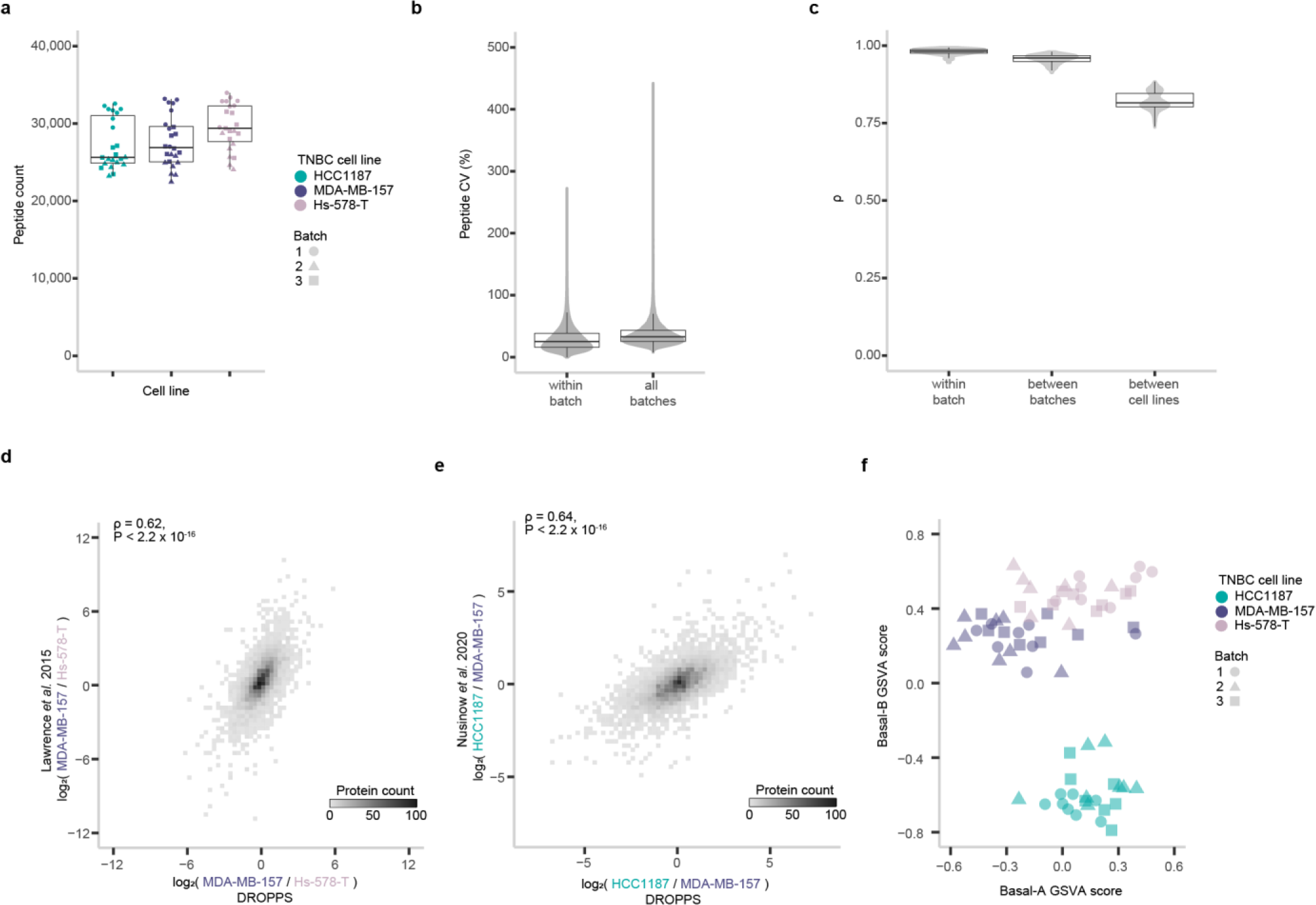
Evaluation of DROPPS for profiling TNBC cell lines differences across batches. (a) The peptide counts of the individual runs. (b) Violin plots of peptide intensity CVs for runs within a batch or for combining batches. (c) Spearman’s ρ between runs of the same cell line within batch, runs of the same cell line between batches, and between all other runs. (d,e) log_2_ fold-changes between cell lines, using data acquired by DROPPS compared to data acquired by (d) Lawrence *et al.*^14^ and (e) Nusinow *et al*.^13^ shown with Spearman’s ρ. (f) GSVA values for Basal A and Basal B TNBC subtypes. Boxplots show the median, interquartile ranges, and 95% confidence interval estimate. TNBC: triple negative breast cancer

**Figure S3.**
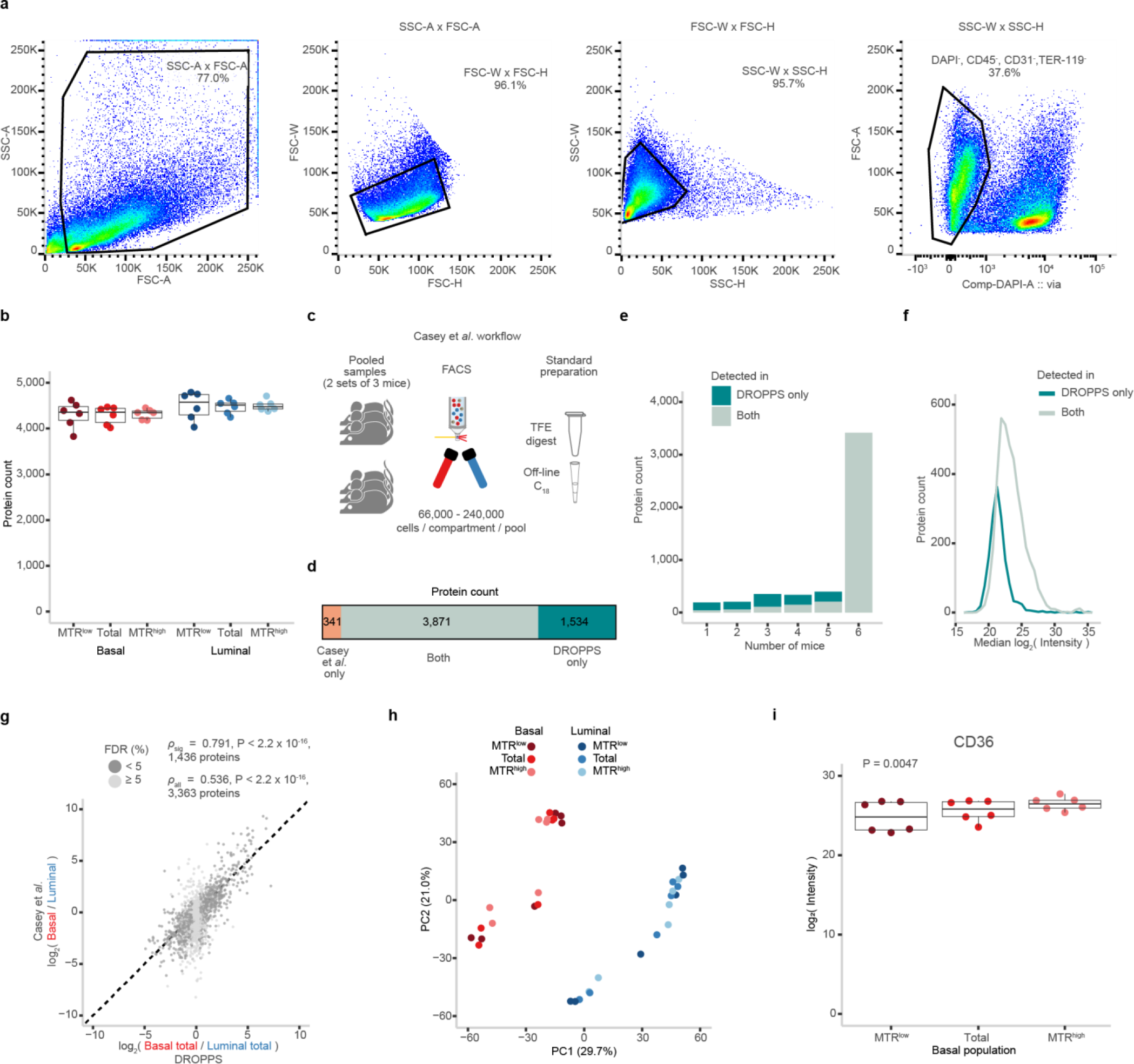
Proteomic profiling of primary mammary epithelial cells sorted by mitochondrial potential. (a) Gating scheme used for FACS purification of epithelial subpopulations. (b) The protein counts of the individual runs shown (*n* = 6 per group) (c) Schematic of experimental workflow used in previous study to sort mammary epithelial compartments. (d) Stacked bar plot showing the protein counts of shared, DROPPS-unique, and Casey *et al.*-unique proteins. (e) Stacked column plot depicting a histogram of the number of individual mice and the set of proteins that were detected using DROPPS including the overlap with our previous dataset (f) Density plot depicting the distribution of log_2_(intensities) for proteins detected using DROPPS and the overlap with our previous dataset. (g) Dot plot depicting the fold change of the total basal and total luminal populations compared to the fold changes from Casey *et al*. where points are colored by significance from the current study. The Spearman’s correlations and significances are shown for all and significantly different proteins along with *y* = *x* for improved visualization. (h) First two principal components of PCA. (i) The log_2_(intensities) of CD36 with repeated measures ANOVA p-value. Boxplots show the median, interquartile ranges, and 95% confidence interval estimate. MTR: MitoTracker Red.

**Figure S4.**
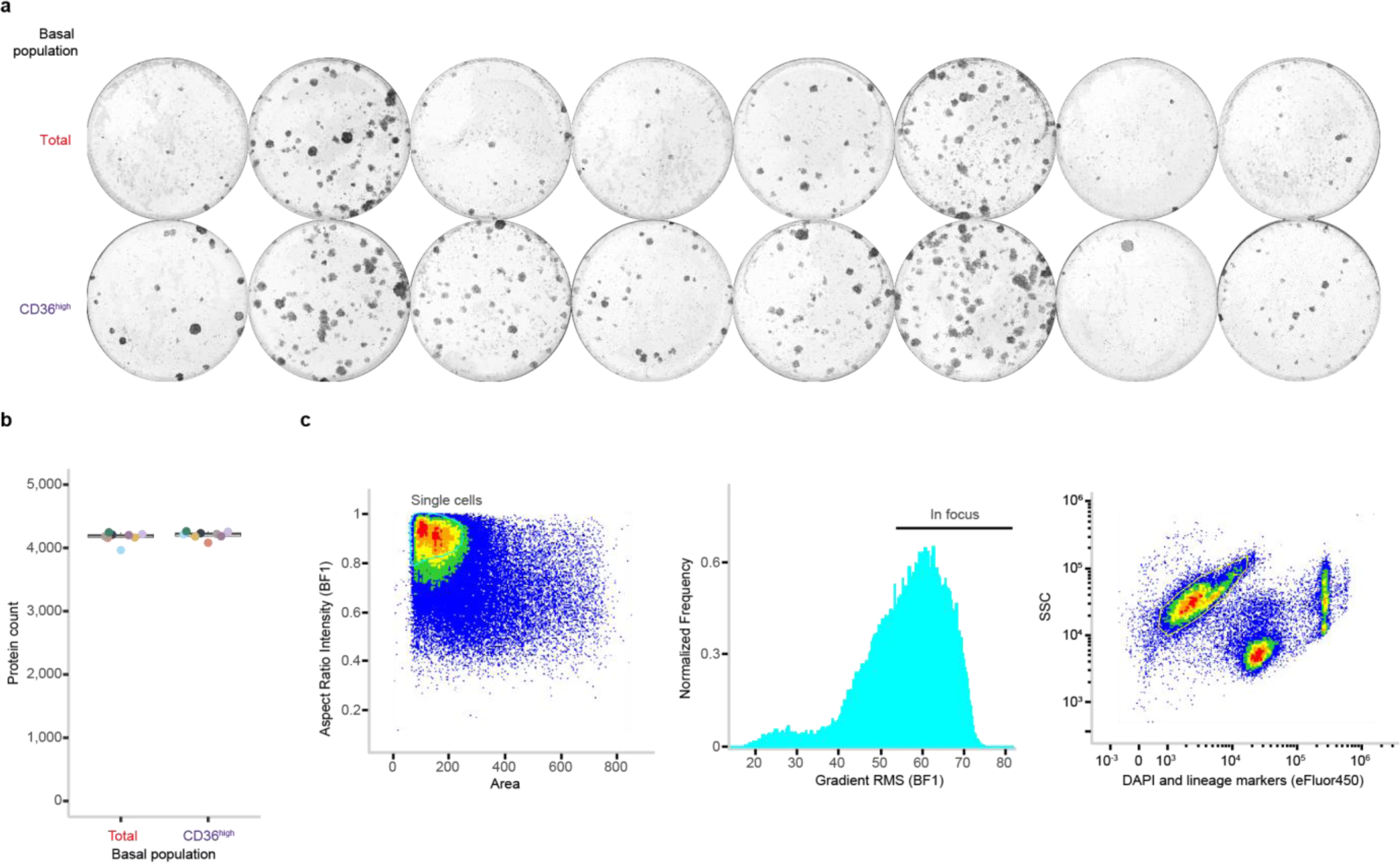
Investigating the phenotype of CD36^high^ basal mammary epithelial cells. (a) CFC assay images of basal mammary epithelial cells sorted by total or CD36^high^ basal populations. (b) The protein counts of the individual runs colored by mouse (*n* = 8). (c) Gating scheme used to analyze mammary epithelial cells by imaging flow cytometry.

## Notes

### Competing Interest Statement

The authors have declared no competing interest.

